# Blood Based Biomarkers of DNA Methylation Associated with Platinum Resistance in High Grade Serous Ovarian Cancer

**DOI:** 10.64898/2026.04.23.720362

**Authors:** Elnaz Abbasi Farid, Shu Zhang, Horacio Cardenas, Zhen Fu, Adam Vieth, Collin M. Coon, Jian-Jun Wei, Daniela Matei, Kenneth P. Nephew

**Affiliations:** Medical Sciences Program, Indiana University School of Medicine-Bloomington, Bloomington, IN 47405, USA; Department of Obstetrics and Gynecology, Feinberg School of Medicine, Northwestern University, Chicago, IL 60611, USA; Bioinformatics and Biostatistics Core, Van Andel Institute, Grand Rapids, MI 49503, USA; Department of Pathology, Feinberg School of Medicine, Northwestern University, Chicago, IL 60611, USA; Robert H Lurie Comprehensive Cancer Center, Chicago, IL 60611, USA; Houston Methodist Neal Cancer Center, Houston, TX, 77030; Department of Anatomy, Cell Biology and Physiology, Indiana University School of Medicine Indianapolis, IN 46202, USA; Indiana University Melvin and Bren Simon Comprehensive Cancer Center, Indianapolis, IN 46202, USA

**Author notes:** Corresponding authors Kenneth P. Nephew, PhD, 302 Biology Building, Indiana University School of Medicine-Bloomington, 1001 E. 3rd St, Bloomington, IN 47405, Daniela Matei, MD, Houston Methodist Neal Cancer Center, 6445 Main Street, Houston, TX, 77030. Equal Contributions (Elnaz Abbasi Farid, Shu Zhang, Horacio Cardenas, Zhen Fu).

**Keywords:** high grade serous ovarian cancer, epigenetics, peripheral blood mononuclear cells, DNA methylation, hypomethylating agent, DNA methyltransferase inhibitor, biomarkers

## Abstract

**Background:** High grade serous ovarian cancer (HGSC) is initially a responsive tumor to platinum (Pt)-based therapy. Pt resistance in HGSC is associated with epigenetic modifications and hypomethylating agents (HMAs) have been studied as carboplatin resensitizing agents. As DNA methylation is detectable in cancer cells and in blood, here we aimed to develop a blood-based methylation signature associated with cancer and cancer recurrence in HGSC.

**Results:** We evaluated genome-wide DNA methylation in de-identified peripheral blood mononuclear cells (PBMCs) from women 1) without cancer (controls, n=20); 2) newly diagnosed HGSC (prior to treatment, Pt-naive, n=60) 3) Pt-resistant recurrent HGSC before and after treatment with the novel HMA/DNA methyltransferase inhibitor (DNMTI) guadecitabine (Pt-resistant, n=30). The Pt-resistant patients were enrolled in NCT02901899 clinical trial testing guadecitabine and the PD-1 inhibitor pembrolizumab. DNA extracted from PBMCs was analyzed by using Infinium MethylationEPIC BeadChips. There were 30,369 differentially methylated loci (DMLs) in Pt-naïve patients vs. controls (adj. p < 0.05, β >10%), with most loci being demethylated. Enriched pathways in PBMCs from cancer patients included *mechanisms of cancer, neutrophil degranulation, and cancer-related signaling pathways (PI3K/AKT, STAT3, HGF, interleukins)*. The number of DMLs was greater (880 DMLs; adj. p<0.05, β>10%) in Pt-resistant vs. Pt-naïve patients, and top enriched pathways associated with Pt-resistant HGSC included pathways in *cancer, metabolic pathways, platelet activation, ABC transporters and signaling pathways (calcium, PI3K/AKT, MAPK, Ras, ErbB, Hippo, Wnt)*. Massive genomewide hypomethylation 5 days after treatment with guadecitabine was observed (13,742 DMLs; adj. p<0.05, β>10%), which persisted 30 days after discontinuation of treatment. Pathways enriched by hypomethylated genes in PBMCs following guadecitabine treatment interestingly included pathways related to n*euronal signaling, such as glutaminergic receptor signaling, axonal guidance signaling, synaptic long-term depression, synaptogenesis signaling and serotonin receptor signaling*. Deconvolution analysis of the methylome data of PBMCs from Pt-resistant recurrent HGSC before versus after HMA treatment predicted *increased* naïve B cells, memory and naïve CD+ T cells, naïve CD4+ T cells, and neutrophils and *decreased* monocytes.

**Conclusions:** We propose new DMLs associated with Pt-naive versus Pt-resistant HGSC. These findings can lead to new biomarkers for HGSC.

## Background

High grade serous ovarian cancer (HGSC), the most common histological type of ovarian cancer, harbors p53 mutations and high initial responsiveness to platinum (Pt)-based chemotherapy(1, 2). However, most tumors relapse, become chemo-resistant and ultimately fatal(2). It has been postulated that HGSC progression to a Pt-resistant state is associated with epigenomic alterations, including increased DNA methylation and altered histone marks deposition (3, 4) leading to transcriptional silencing of tumor suppressor (TSGs) and other pro-apoptotic genes(5, 6). Tumor epigenetic signatures have been associated with clinical outcomes (7) (8, 9) highlighting clinical significance and potential for therapeutic targeting. DNA methylation and histone modifications are intimately linked (10, 11) and regulate the assembly of transcriptionally permissive or repressive (i.e., open or closed) chromatin. The concept that DNA methylation and histone modifications are associated with chemoresistance in HGSC has been proposed by us and others (11-13).

Hypomethylating agents (HMAs) have been shown to restore Pt sensitivity in chemo-resistant OC cell lines and xenografts (14) (15-19) (20-22). Moreover, clinical trials in women with Pt-resistant HGSC using DNA-methylation transferase inhibitors (DNMTI) prior to Pt in a chemo-resensitization strategy reported significant clinical activity (23, 24). Furthermore, CpG island methylation in tumor biopsies obtained before and after treatment showed that the number of demethylated genes was greater in tumor biopsies from patients achieving improved disease control (23, 24). Treatment-induced hypomethylation of TSGs and corresponding gene expression in paired tumor biopsies was also reported, and the findings pointed to a potential panel of genes epigenetically regulated and mechanistically associated with Pt resistance (24) and thus proposing new potential methylation biomarkers (23, 24).

As obtaining tumor biopsies from patients on clinical trials remains difficult, DNA methylation profiling of cell-free DNA (cfDNA) in the circulation or in peripheral blood mononuclear cells (PBMCs) has recently emerged as a potential non-invasive liquid biopsy tool for cancer patients. Previous studies have associated methylation changes in PBMCs with tumor progression in various cancers, including breast, colorectal, ovarian and head and neck cancers (25), suggesting potential utility in early detection or monitoring cancer progression (26). Unlike cfDNA or circulating tumor cells (CTCs), which are present in low amounts, PBMCs are abundant, easier to extract and offer longer DNA stability. In addition, analysis of PBMCs represents a highly repeatable, minimally invasive approach with broad applicability (27), rendering it highly suitable for retrospective and prospective analyses. Furthermore, PBMC methylation changes have the potential to serve as biomarkers for response to epigenetic treatments in cancer patients (11). However, the effect of DNMTIs on PBMCs from HGSC patients remains to be established.

In the current study, we analyzed methylome changes in PBMCs collected from control patients (without cancer), women with newly diagnosed HGSC, and women treated on our recent clinical trial targeting DNA methylation with the HMA guadecitabine to improve response to immunotherapy (28). We sought to detect differences in PBMCs methylomes in women with cancer compared to controls and in PBMCs from women with Pt-resistant versus Pt-naive HGSC. Furthermore, we sought to define the effects of the HMA studied in our trial on PBMCs methylomes and identify cancer relevant genes and pathways disturbed by the inhibitor. Here we report the analysis of the PBMC methylomes in the above contexts. Distinct blood-based DNA methylation patterns differentiated Pt-naive from non-cancer and Pt-resistant HGSC. Exposure of PBMCs to DNMTI resulted in profound hypomethylation of genes and pathways relevant to cancer processes in HGSC, as well as changes in peripheral immune cell composition before and after HMA treatment. Our findings support the potential of blood-based DNA methylation as a non-invasive biomarker for patient response to HMA.

## Materials and Methods

### Human specimens

PBMCs were collected from platinum (Pt)-naive (at diagnosis) and from Pt-resistant (enrolled in clinical trial NCT02901899) HGSC patients. The study protocol was approved by Institutional Review Boards (IRB) to protect participant safety and maintain compliance with the Declaration of Helsinki. All patients provided informed consent for tumor and blood banking for research analyses. PBMCs were isolated as follows: 1) 60 Pt-naive samples obtained at diagnosis of HGSC from the Pathology Core at Lurie Cancer Center; 2) Pt-resistant samples collected at baseline (cycle 1, day 1; C1D1), before administration of guadecitabine and pembrolizumab; 3) then after treatment on C1D5 (patients received guadecitabine on Days 1-4 and pembrolizumab on Day 5 (30 paired C1D1 and C1D5 samples); 4) de-identified female subjects without cancer served as “normal” controls (n=20; Zen-Bio, Inc., NC, USA).

### DNA extraction

DNA extraction from PBMCs was performed by using the DNeasy Blood & Tissue Kit (Qiagen Sciences Inc., Maryland, USA, Cat 69506), according to the manufacturer’s protocol. The DNA quality and quantity were assessed using NanoDrop 2000 spectrophotometer (Thermo Fisher Scientific, Massachusetts, USA) for purity estimates, and Qubit Fluorometer (Thermo Fisher Scientific, Massachusetts, USA) to measure DNA concentration. All samples yielded high-quality DNA suitable for downstream analyses. To differentiate between methylated and unmethylated cytosines, DNA underwent bisulfite conversion using a kit (Zymo Research, California, USA, Cat. D5031) and following the manufacture’s protocol.

## Methylation studies

Genome-wide methylation profiling was conducted using the Infinium HumanMethylationEPIC v2.0 BeadChip platform (Illumina, California, USA). This platform detects methylation levels at over 935,000 CpG sites. The process included DNA hybridization to bead arrays, enzymatic extension, and fluorescent staining, enabling precise measurement of methylation at individual sites.

### Bioinformatic Analysis

The raw methylation data were processed with the SeSAMe (Signal Extraction and Summarization of Array Methylation Experiments, v1.24.0) R package (29, 30). This software normalized signal intensities, corrected for dye bias, and filtered low-quality probes, ensuring the reliability of the resulting β-value matrices. These β-values, which range from 0 (unmethylated) to 1 (fully methylated), were used to identify differentially methylated loci (DMLs) and differentially methylated regions (DMRs) between groups. A β-value threshold of 0.1, along with an adjusted P-value of <0.05, was applied to identify significant changes, with the Benjamini-Hochberg method ensuring control over the false discovery rate.

To provide additional context, CpG sites were annotated based on genomic location relative to CpG islands, shores, shelves, and transcription start sites (TSS). CpG islands were defined as regions of DNA greater than 500 bases long, with a GC content exceeding 55% and a high observed-to-expected CpG ratio. Regions extending up to 2 kilobases (kb) upstream or downstream of these islands were labeled as CpG shores, while areas 2–4 kb away were defined as CpG shelves. Sites outside these regions were classified as “open sea.” In relation to gene transcripts, CpG sites were further categorized based on their proximity to key genomic features. For example, sites within 200 bases upstream of a transcription start site were classified as TSS200, while those between 200 and 1500 bases upstream were labeled as TSS1500.

Additional annotations included sites in the 5′ untranslated region (5′UTR), the first exon, and gene body. CpG sites without any of these associations were described as “not linked to a gene.” CpG site annotation information was obtained from sesameData R package (v1.24.0) (30)and Noguera-Castells et al. (31). We also explored the methylation patterns of CpG sites in the Long Interspersed Nuclear Element-1 (LINE-1) regions. To do this, we downloaded RepeatMasker tracks for the hg38 genome from the UCSC Genome Browser and used the GenomicRanges R package (v1.58.0) (32) to identify overlapping probes in EPIC v2 located within LINE-1 regions.

### Pathway Analyses

To define pathways and identify functional interactions of genes differentially methylated between groups, the following approaches were used: 1) Ingenuity Pathway Analysis (IPA) (QIAGEN Digital Insights, California, USA), a comprehensive database for curated molecular pathways and upstream regulator analysis; 2) KEGG, which maps genes to metabolic and signaling pathways to understand cellular processes; 3) GO, which categorizes genes by biological processes, molecular functions, and cellular components; 4) TF analysis, which identifies regulatory proteins potentially driving methylation changes and 5) WP, a collaborative platform providing additional annotation and validation of pathways.

### Deconvolution Analysis

To determine immune cell composition, deconvolution analysis of the methylation data was conducted by using R package EpiDISH (v2.22.0) (33) to infer the fractions of a priori known cell subtype present. We first used the “mLiftOver” function from SeSAMe to convert the EPIC v2 to EPIC v1 format, as the reference used in epiDISH did not natively support EPIC V2. Next, in the deconvolution analysis, we used “epidish” function with the Robust Partial Correlations (RPC) method (34), using cent12CT.m as the reference data. This reference included 12 immune cell types: naive and mature B-cells, naive and mature CD4T-cells, naive and mature B-cells, T-regulatory cells, NK-cells, neutrophils, monocytes, eosinophils, and basophils. Next, we used the R package glmmTMB (35) to perform a beta mixed-effects regression analysis on cell type percentages between groups of samples.

Additionally, we used the R package emmeans (36) to estimate means and p-values between sample groups.

### Data availability

The data generated during this study are publicly available in the Gene Expression Omnibus (GEO) repository, hosted by the National Center for Biotechnology Information (NCBI), under the accession number GSE315041.

## Results

### PBMC methylomes in Pt-naive HGSC compared to non-cancer

To begin to characterize DNA methylation patterns of PBMCs, we first compared methylomes from non-cancer (“normal”) PBMCs to samples from newly diagnosed HGSC (Pt-naive). Principal Component Analysis (PCA) of methylation profiles revealed distinct clustering between Pt-naive and non-cancer PBMCs (**Fig. 1A**). PC1 and PC2 explained 42.64% and 9.05% of the variance, respectively, capturing most of the variability in the dataset. Pt-naive PBMCs exhibited greater dispersion, indicating greater heterogeneity, while non-cancer PBMCs clustered more tightly, reflecting consistent methylation profiles. Results of differential methylation between non-cancer and Pt-naive HGSC are shown as volcano plots (**Fig. 1B**). The number of differentially methylated loci (DMLs) between non-cancer versus Pt-naive HGSC PBMCs was significant (adj. p<0.05, β >10%), and the majority of the 30,369 DMLs were demethylated, which is indicated by the shift to the left of the red dots representing differentially methylated CpG sites (**Fig. 1B**). We then examined the pathways enriched by the hypomethylated genes. Canonical pathway analysis of the DML in non-cancer versus Pt-naive PBMCs revealed the top seven most significantly altered pathways (**Fig. 1C**), all of which were cancer-related, including *mechanisms of cancer, neutrophil degranulation, and cancer-related signaling pathways (PI3K/AKT, STAT3, HGF, interleukins)*. Overall, up- and down-regulated pathways and molecules influenced by hypomethylated genes in Pt-naive HGSC versus non-cancer PBMCs are illustrated (**Fig. 1D**; orange (upregulated) vs. blue (downregulated)).

**Figure 1.**
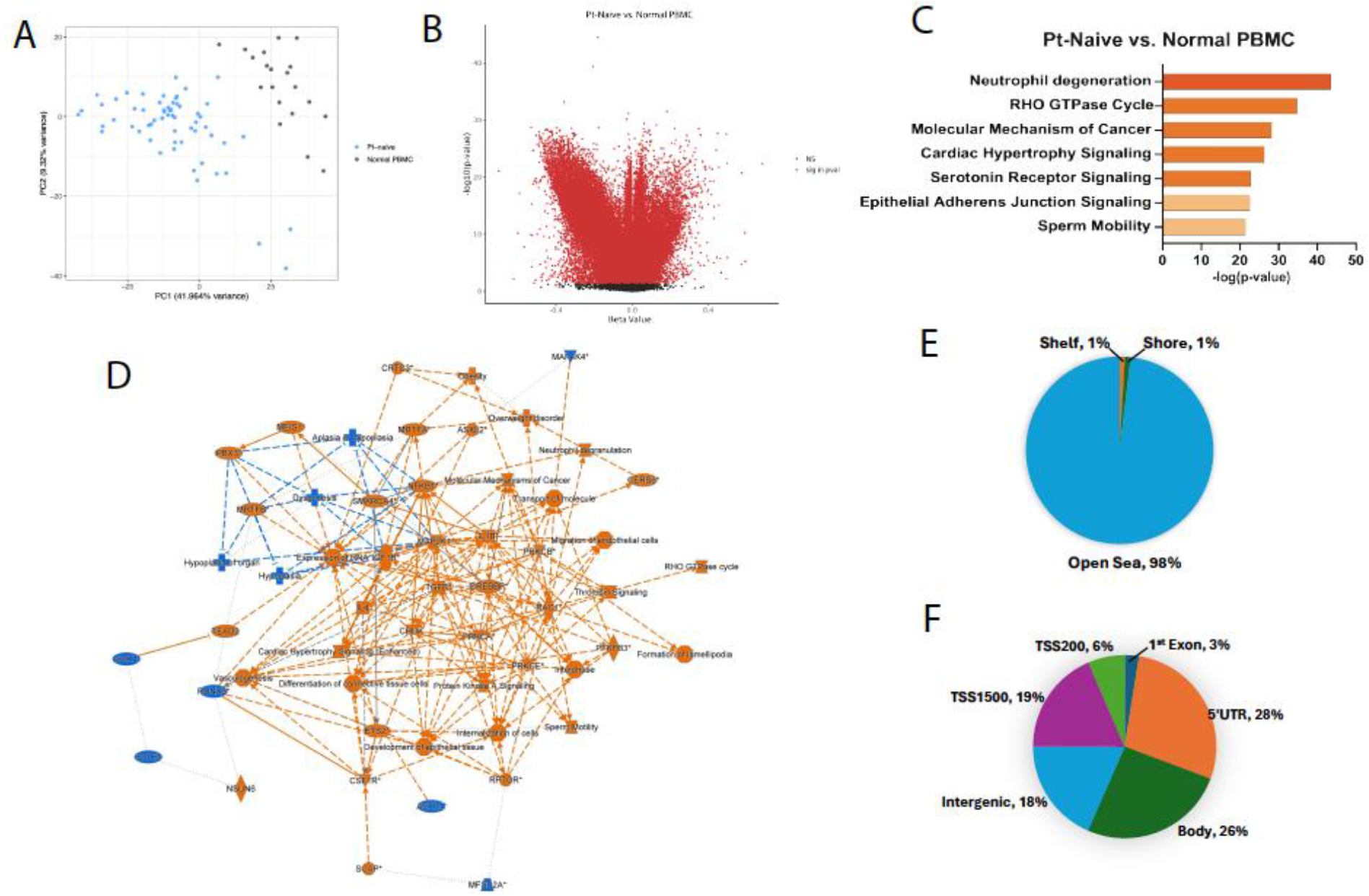
Comparison of PBMC methylation from non-cancer controls compared to patients with platinum-naive HGSC. (A) Principal Component (PC) analysis of PBMCs from Pt-naive vs. normal PBMCs; (B) Volcano plot showing differentially methylated loci (DMLs) between Pt-naive vs. normal PBMCs; (C) Canonical pathway analysis highlighting the top seven most significantly altered pathways; (D) Overall upregulated and downregulated pathways in Pt-naive vs. normal PBMCs, (E, F) Pie chart showing the genomic distribution of differentially methylated CpG sites relative to CpG island or a gene.

The CpG probes interrogated on the Infinium HumanMethylationEPIC BeadChip were distributed between CpG islands, regions flanking the CGIs (within 4Kb of the nearest island) referred to as “shelves” and “shores” and representing another third of all sites, and sites unrelated to an island, generally occurring within the gene body and termed as “open sea” sites, which represent another third of all probes. The genomic distribution of differentially methylated CpG sites relative to a CpG islands or a gene was markedly different between the two groups. In non-cancer compared to Pt-naive PBMCs, essentially all (98%) DMLs resided in the open sea, with 0-1% in the other genomic locations (island, shores, shelves; **Fig. 1E**). Relative to genes, of the total hypomethylated sites, 28% were within 1,500 bp of the TSS or in the first exon, whereas another 28% and 26%, respectively were in 5’ UTR and gene bodies, followed by 18% in intergenic regions (**Fig. 1F**).

### PBMC methylomes in platinum-resistant compared platinum-naive HGSC patients

Next, we analyzed PBMC methylomes from Pt-naive compared to Pt-resistant HGSC patients. The two groups displayed distinctly different clusters based on PCA analysis (**Fig. 2A**). PC1 and PC2 captured most of the variability in the dataset, 44.26% and 10,84% of the variance, respectively. Pt-naive PBMCs clustered more tightly, reflecting consistent methylation profiles. The observed greater dispersion of methylation profiles of PBMCs from Pt-resistant patients indicated greater PBMC methylome heterogeneity. As shown in the volcano plot in **Fig. 2B**, the number of DMLs in Pt-resistant PBMCs was greater compared to Pt-naive (30,369 DMLs; adj. p<0.05, β >10%). Canonical pathway analysis of the DML in Pt-naive vs. Pt-resistant PBMCs revealed that the top nine most significantly altered pathways were enriched in cancer-related functional processes and signaling pathways, such as *glioblastoma signaling, pre-Notch processing, and bladder cancer signaling* (**Fig. 2C**). Inhibition of key pathways related to *differentiation of phagocytes, myeloid leukocytes, hematological progenitors, antigen presenting cells, homing of cells, blood vessel formation* and others was predicted in Pt-naive vs. Pt-resistant PBMCs (**Fig. 2D**; blue vs. orange). Together, the DMLs between Pt-naïve and Pt-resistant PBMC affect predominantly pathways related to immune cell differentiation suggesting inhibition of these processes in PBMCs from patients with more advanced cancer. The genomic distribution of differentially methylated CpG indicated that the vast majority (81%) the DMLs resided in the open sea, with 19% in other genomic locations (island, shores, shelves; **Fig. 2E**). Relative to genes, of the total DMLs, 26% were within 1,500 bp of the TSS or in the first exon, whereas another 27% and 24%, respectively were in 5’ UTR and gene bodies, and 23% in intergenic regions (**Fig.2F**).

**Figure 2:**
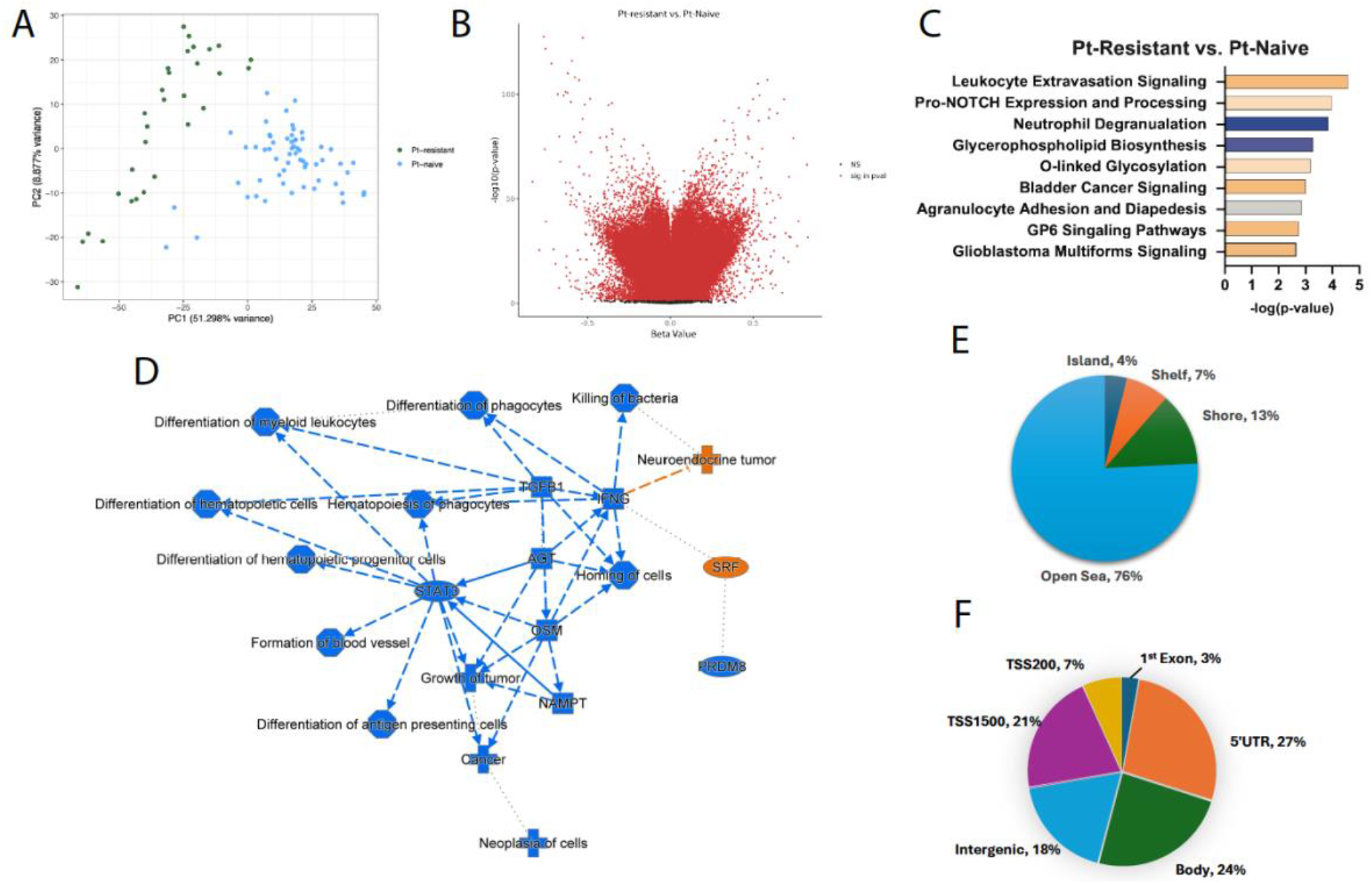
Comparison of PBMC methylation from patients with platinum-resistant HGSC compared to platinum-naïve HGSC. (A) Principal Component (PC) analysis of PBMCs from Pt-naive vs. Pt-resistant; (B) Volcano plot showing differentially methylated loci (DMLs) between Pt-naive vs. Pt-resistant, (C) Canonical pathway analysis highlighting the top nine most significantly altered pathways; (D) Overall upregulated and downregulated pathways in Pt-naive vs. Pt-resistant; (E,F) Pie chart showing the genomic distribution of differentially methylated CpG sites relative to CpG island or a gene.

### Genome-wide effects of HMAs on the PBMC methylome from HGSC patients

Although HMAs have been shown to induce hypomethylation directly in HGSC tumors (8, 9, 24), an extensive analysis of the global impact of HMA exposure on the PBMCs in HGSC patients has not been conducted. PCA analysis of PBMCs from patients with Pt-resistant HGSC (C1D1), and guadecitabine-treated (C1D5), indicated that the groups were intermixed with no distinct clusters (**Fig. 3A**), suggesting marked heterogeneity among the PBMC methylomes. PC1 and PC2 captured most of the variability in the dataset, 44.26% and 10,84% of the variance, respectively. We observed a dramatic shift to the left of the red dots representing differentially methylated CpG sites (**Fig. 3B**), demonstrating exposure of PBMCs to guadecitabine had a profound demethylation effect (13,742 DMLs; adj. p<0.05, β >10%). Canonical pathway analysis of DMLs revealed top nine most significantly altered pathways were highly enriched for various signaling pathways in HMA-treated versus Pt-resistant baseline PBMCs, including pathways related to neuronal signaling, such as *glutaminergic receptor signaling, axonal guidance signaling, synaptic long-term depression, synaptogenesis signaling and serotonin receptor signaling* (**Fig. 3C**). Activation of neuronal signaling has been reported as a driver of cancer progression, including recent reports in HGSC (37-39). Predicted activation of key growth cancer-associated pathways among DMLs from HMA-treated versus Pt-resistant baseline PBMCs is shown (**Fig. 3D**; orange denotes upregulated vs. blue downregulated). Furthermore, differences in the distribution of the differentially methylated CpG sites was observed between the two groups. In guadecitabine-treated compared to Pt-resistant baseline samples, most of the DMLs were found in the open sea (81%) versus islands (1%) and shores and shelves each with 9% (**Fig. 3E**). Of all the hypomethylated sites in guadecitabine-treated versus Pt-resistant PBMCs, 26% were within 1,500 bp of the TSS or in the first exon of a gene, whereas another 27% and 24% were found in the 5’ UTR and gene bodies, respectively, followed by 23% in intergenic regions (**Fig. 3F**).

**Figure 3:**
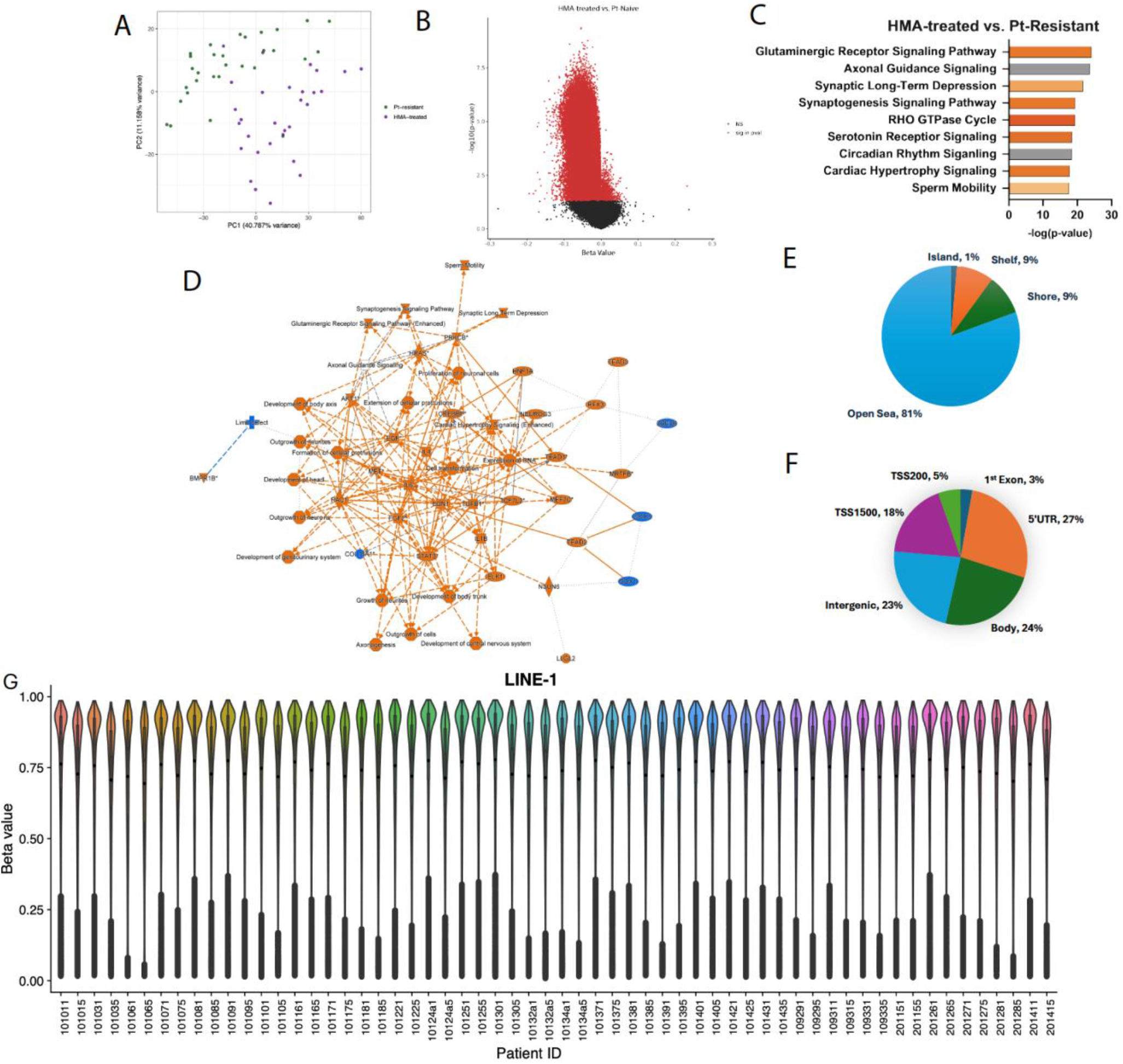
Comparison of PBMC methylation from platinum-resistant compared to HMA-treated HGSC patients. (A) Principal Component (PC) analysis of PBMCs from Pt-resistant vs. HMA-treated HGSC; (B) Volcano plot showing differentially methylated loci (DMLs) between Pt-resistant vs. HMA-treated; (C) Canonical pathway analysis highlighting the top nine most significantly altered pathways; (D) Overall upregulated and downregulated pathways in Pt-resistant vs. hypomethylating agent-treated; (E,F) Pie chart showing the genomic distribution of differentially methylated CpG sites relative to CpG island or a gene. (G) Violin plot of beta values distribution of LINE-1 CpG sites in each PBMC sample. Each violin represents one sample before and after HMA treatment. The last number “1” indicates before treatment, and “5” indicates post treatment.

Global DNA methylation of long interspersed nuclear element-1 (LINE-1) was subsequently analyzed between Pt-resistant and HMA-treated PBMC. Violin plots analysis of CpG site methylation levels across paired patient samples demonstrated consistent bimodal distributions, with shifts in intermediate β-value ranges towards lower β-value in PBMCs from HMA treated patients (**Fig. 3G**). Hierarchical clustering and heatmap of LINE-1 CpGs further indicated a clear segregation of profiles between PBMCs obtained before versus after HMA treatment (**Supplementary Fig. S1**).

### Predicted immune cell composition

As HMAs such as azacitidine and decitabine induce global DNA demethylation in PBMCs, it was of interest to predict the composition and proportions of different immune cell types induced by guadecitabine based on methylomic profiles. To predict and compare the cellular make-up of the PBMCs, deconvolution analysis of the methylome data of PBMCs from Pt-resistant recurrent HGSC before versus after HMA treatment was performed (**Fig. 4A, B**). This analysis predicted a significant *increase* naïve B cells, memory and naïve CD+ T cells, naïve CD4+ T cells, and neutrophils and a significant *decrease* in monocytes induced by HMA treatment (**Fig. 4C**). Other immune cell type populations were not predicted to be significantly altered (**Supplemental Fig. S2**).

**Figure 4.**
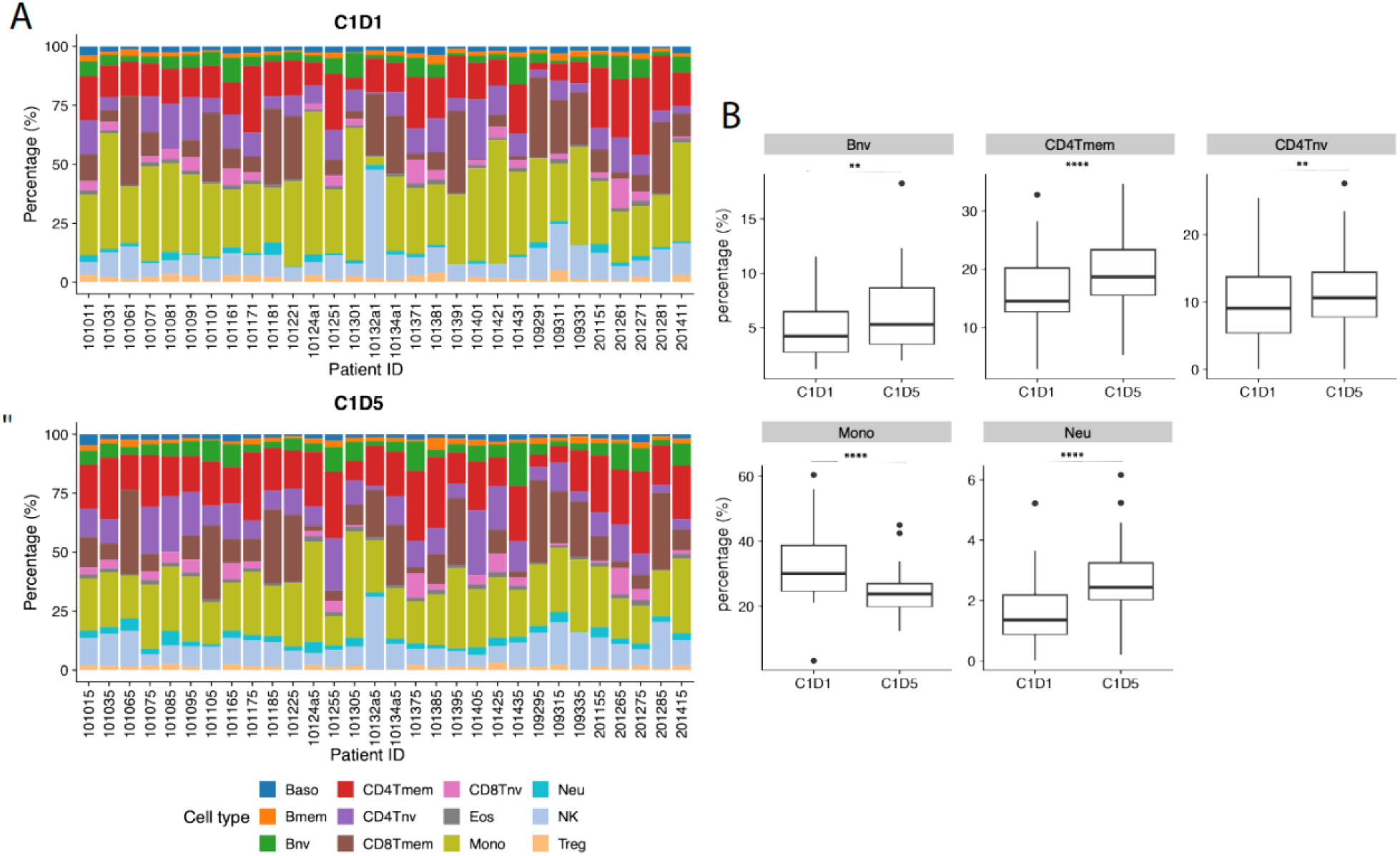
Peripheral immune cell composition before and after HMA treatment. (A-B) Stacked bar plots display the relative proportions of major immune cell types in peripheral blood mononuclear cell (PBMC) samples from before HMA treatment (C1D1) and after treatment (C1D5), as estimated by methylation-based deconvolution. (C) Boxplots depicting the percentage (%) of five statistically significant peripheral immune cell subpopulations: naive B cells (Bnv), memory CD4+ T cells (CD4Tmem), naive CD4+ T cells (CD4Tnv), monocytes (Mono), and neutrophils (Neu) at before (C1D1) and after (C1D5) HMA treatment.

## Discussion

This study demonstrates that genome wide DNA methylation is measurable in PBMCs, distinguishing healthy individuals from patients with HGSC pointing to a distinct cancer associated signature. Importantly, we also detect clear differences in the PBMC methylome of patients with Pt-naïve compared to Pt-resistant HGSC, indicating that prior cancer therapy leaves a lasting imprint on peripheral immune cells. In addition, we show that the PBMC methylome is highly altered by treatment with an HMA in the context of a clinical trial testing an HMA with immune checkpoint inhibitor in patients with recurrent HGSC. The detection of disease-associated methylation patterns in circulating immune cells highlights the potential systemic epigenetic perturbations that accompany tumor development and tumor evolution through treatment supporting feasibility of blood-based epigenomic profiling as a non-invasive cancer detection or cancer monitoring strategy. Our study has several implications.

First, we show significant differences in the methylome of PBMCs from patients with HGSC versus healthy women, with over 30,000 DMLs detected, of which most were hypomethylated. Differences in PBMC methylation patterns associated with cancer have been recently reported for breast, colorectal and lung cancer patients (40, 41). In addition, differences in methylation of specific CpGs have been associated with clinical parameters, such as overall survival in a recent lung cancer study (42). Together with the present findings, these studies highlight the potential of developing PBMC DNA methylation as a diagnostic biomarker, either alone or in combination with DNA methylation patterns from cfDNA or tumor biopsies.

Second, PBMC methylation profiles differentiated Pt-naïve from Pt-resistant HGSC, indicating that treatment history and the emergence of chemoresistance are reflected in distinct, measurable epigenetic states of the immune compartment. This could reflect direct chemotherapy-induced altered DNA methylation due to Pt-induced DNA damage (43), or alterations in the PBMC composition associated with DNA methylation changes during the evolution of the disease. In support of the first theory, Flanagan et al. reported that Pt-based treatment causes specific CpG methylation changes in blood, because of chemotherapy-induced functional mismatch repair (43). Interestingly, they observed similar methylation changes in both blood and tumors at the time of relapse (43). In our study, we did not have access to paired PBMC and tumor specimens, limiting the strength of our conclusions, but in a previous study using tumor samples, we also reported significant differences in CpG site methylation in tumors at diagnosis vs. Pt-resistant recurrence (9). Furthermore, pathway analyses of PBMC DMLs revealed that key cancer signaling networks were altered along with emergence of Pt-resistance, along with inhibition of leukocytes and phagocyte differentiation and other immune functions. These findings are also consistent with our previous observation from analyses of tumor specimens (9), suggesting that systemic immune epigenetic remodeling may accompany or contribute to the development of Pt-resistance and providing a new dimension through which to understand the biology of refractory HGSC.

Third, we show that treatment with a guadecitabine-based regimen resulted in robust and persistent genome-wide hypomethylation in PBMCs, consistent with the known mechanism of action of HMAs. This work builds on our group’s prior investigator-initiated clinical trials evaluating HMAs to re-sensitize ovarian tumors to Pt-based chemotherapy. Across those studies, we showed that HMA exposure induces profound hypomethylation in tumors, and that these changes are detectable in matched tumor biopsies obtained before and after therapy (23, 24, 44, 45). The current PBMC-based findings extend those observations, demonstrating that systemic epigenetic responses mirror tumor-intrinsic changes and could serve as a more accessible surrogate for tumor sampling. Indeed, the marked hypomethylation detected in PBMCs sampled after guadecitabine treatment mirrors the significant decrease in LINE1 methylation measured by pyrosequencing in the same PBMC specimens previously reported and parallels changes in tumor DNA methylation from the same patients (28). In the current study, we observe marked LINE-1 hypomethylation in post-treatment PBMC samples, confirming drug response in the context of successful HMA-induced global DNA demethylation. Furthermore, the HMA-induced demethylation of LINE-1 elements suggests the potential impact on genomic stability and retrotransposon activity (46, 47). Interestingly, others have also reported increased methylation of LINE1 elements in PBMCs from patients with HGSC versus matched controls, although LINE1 methylation was discrepant between paired tumors and PBMCs in that study (48).

Cell-type deconvolution is a bioinformatic method that enables inference of the relative abundance of various cell types from methylation profiles. Deconvolution analyses demonstrated an increased naïve B cell, memory and naïve CD+ T cells, naïve CD4+ T cells, and neutrophils and a corresponding decrease in monocytes. These immune shifts imply that HMAs not only act directly on tumor and immune cell methylomes but also reshape the circulating immune landscape, potentially influencing anti-tumor immunity and treatment responsiveness. Direct effects of this regimen on immune cell populations were also assessed by multi-parameter flow analysis and was previously reported (28). We cannot exclude that some of the observed effects are due to pembrolizumab, which was part of the regimen tested in this trial (28). The lack of transcriptomic data from PBMCs limits functional validation of how the observed methylome changes impacted gene expression in specific immune cells. As the PBMCs analyzed here were not sorted in immune cell subpopulations, we are not able to determine mechanistically how some immune cell types responded to HMA while others did not.

## Conclusions

- Our findings support the development of blood-based DNA methylation profiling as a minimally invasive biomarker for monitoring response to HMAs.
- The ability to detect global and pathway-specific methylation changes in PBMCs provides a practical tool for assessing pharmacodynamic effects, immune modulation, and potentially predicting clinical benefit.
- These results highlight the promise of integrating blood-based methylation biomarkers into clinical trials of epigenetic therapy.
- Together with emerging technologies that measure epigenetic changes in cell free DNA (49-51), methylomic analysis of PBMCs provides direct monitoring of treatment effects in cancer patients. Such approaches may improve patient selection and enable real-time response assessment in patients receiving HMAs.

## Acknowledgments

We thank Marie Adams (Genomics Core, Van Andel Institute, Grand Rapids, MI) and the Jerry W. and Peggy S. Throgmartin Chair in Oncology. This research was funded in in part by Department of Defense Ovarian Cancer Research Program Grant Number OC200078 W81XWH-21-1-0281 and DoD Award W81XWH-17-1-0141--OC160260, P30CA082709-25 and the. The indicated Stand Up To Cancer Grant is administered by AACR, the scientific partner of Stand Up To Cancer. The clinical trial was supported by Merck and Astex, Inc. PBMC from patients with ovarian cancer were banked and retrieved from the Pathology Core supported by NCI CCSG P30CA060553 awarded to the Robert H. Lurie Comprehensive Cancer Center. K.P.N. holds the Jerry W. and Peggy S. Throgmartin Chair in Oncology.

