## Supplementary figures and images for "Blood Based Biomarkers of DNA Methylation Associated with Platinum Resistance in High Grade Serous Ovarian Cancer"

### Supplementary Figure 1-edited.tif

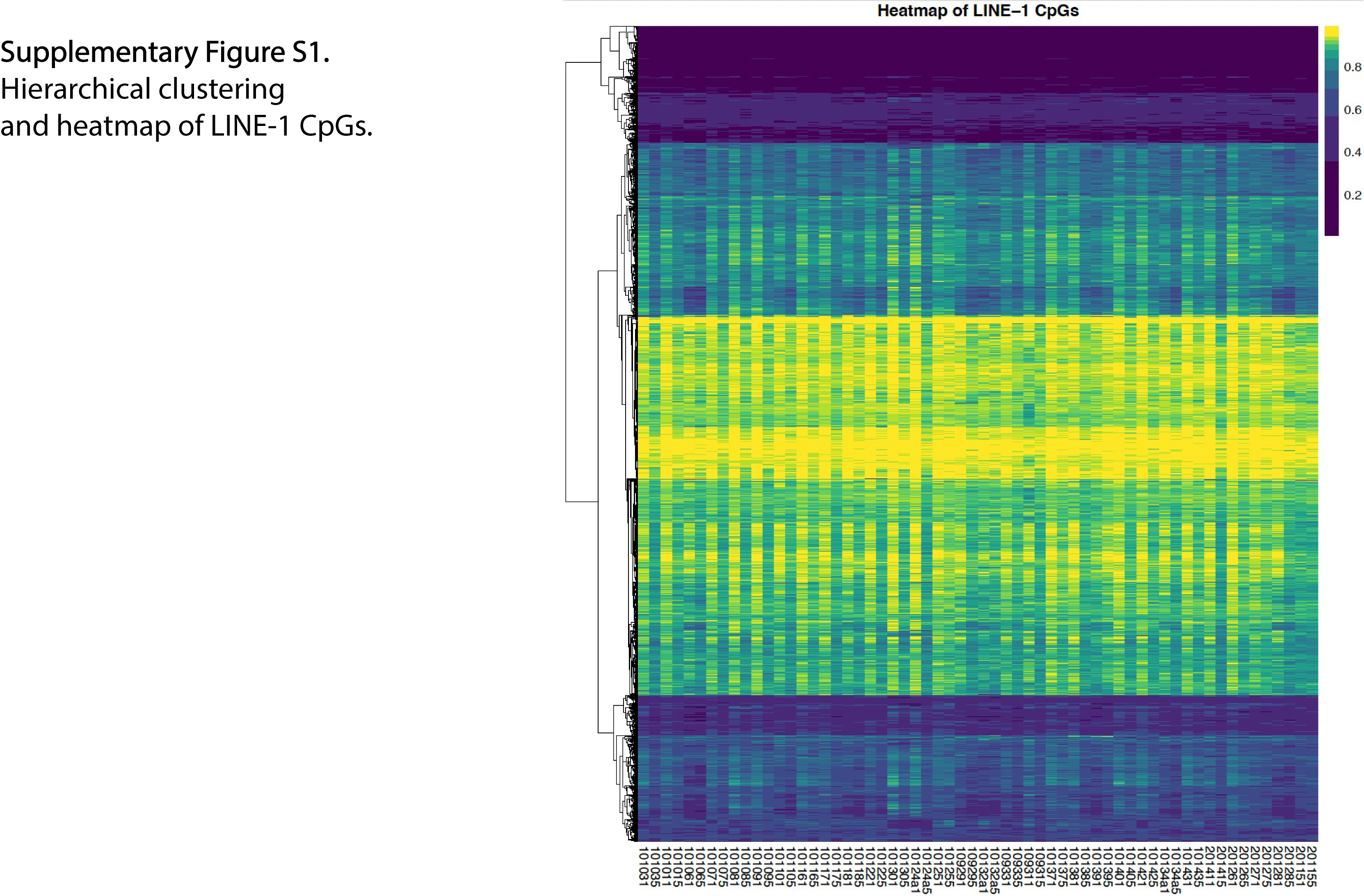

### Supplementary Figure 2-edited.tif

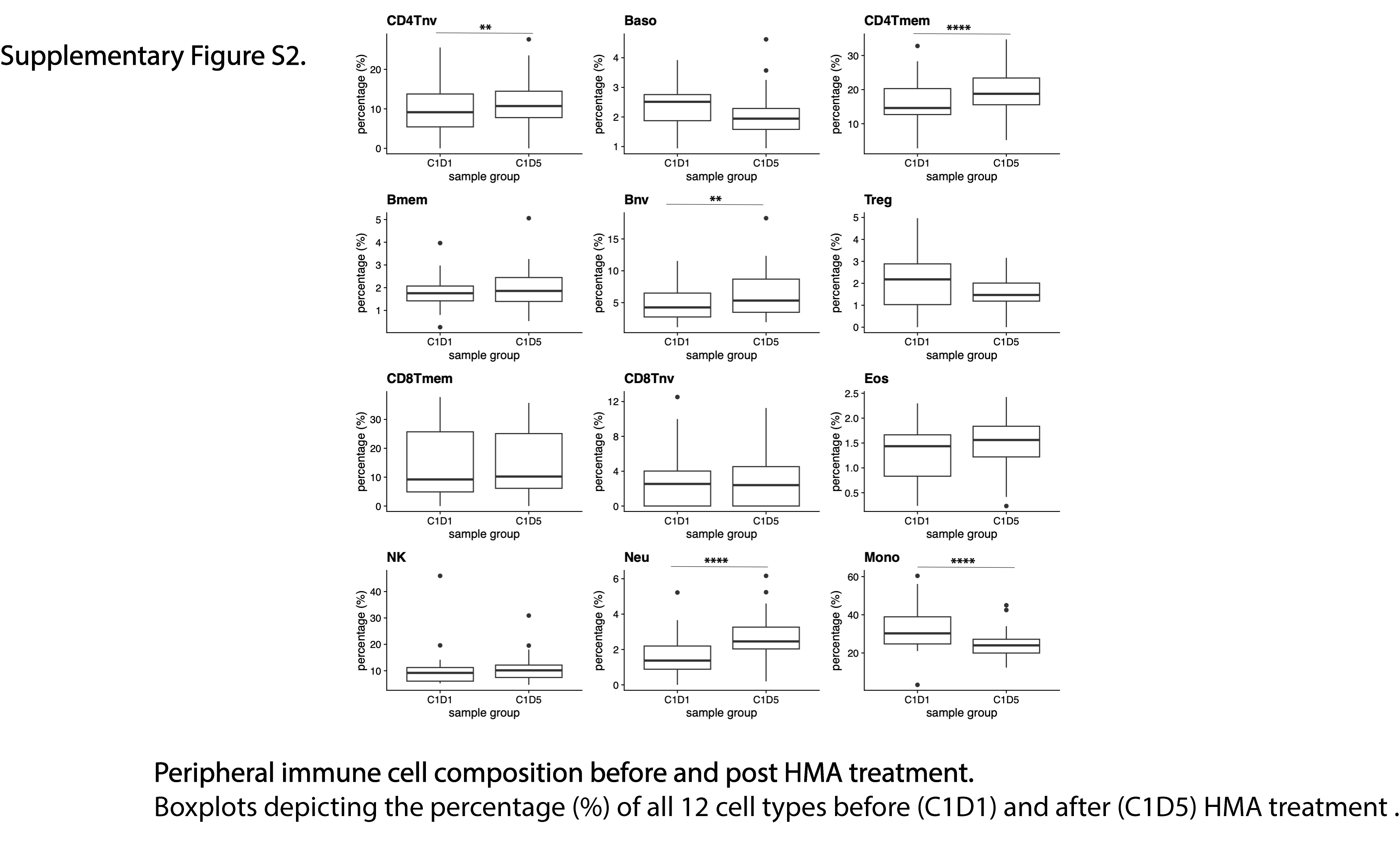
